# Thousands of primer-free, high-quality, full-length SSU rRNA sequences from all domains of life

**DOI:** 10.1101/070771

**Authors:** Søren M. Karst, Morten S. Dueholm, Simon J. McIlroy, Rasmus H. Kirkegaard, Per H. Nielsen, Mads Albertsen

## Abstract

Ribosomal RNA (rRNA) genes are the consensus marker for determination of microbial diversity on the planet, invaluable in studies of evolution and, for the past decade, high-throughput sequencing of variable regions of ribosomal RNA genes has become the backbone of most microbial ecology studies. However, the underlying reference databases of full-length rRNA gene sequences are underpopulated, ecosystem skewed^1^, and subject to primer bias^2^, which hamper our ability to study the true diversity of ecosystems. Here we present an approach that combines reverse transcription of full-length small subunit (SSU) rRNA genes and synthetic long read sequencing by molecular tagging, to generate primer-free, full-length SSU rRNA gene sequences from all domains of life, with a median raw error rate of 0.17%. We generated thousands of full-length SSU rRNA sequences from five well-studied ecosystems (soil, human gut, fresh water, anaerobic digestion, and activated sludge) and obtained sequences covering all domains of life and the majority of all described phyla. Interestingly, 30% of all bacterial operational taxonomic units were novel, compared to the SILVA database (less than 97% similarity). For the Eukaryotes, the novelty was even larger with 63% of all OTUs representing novel taxa. In addition, 15% of the 18S rRNA OTUs were highly novel sequences with less than 80% similarity to the databases. The generation of primer-free full-length SSU rRNA sequences enabled eco-system specific estimation of primer-bias and, especially for eukaryotes, showed a dramatic discrepancy between the *in-silico* evaluation and primer-free data generated in this study. The large amount of novel sequences obtained here reaffirms that there is still vast, untapped microbial diversity lacking representatives in the SSU rRNA databases and that there might be more than millions after all^1, 3^. With our new approach, it is possible to readily expand the rRNA databases by orders of magnitude within a short timeframe. This will, for the first time, enable a broad census of the tree of life.

---

To obtain primer-free and full-length SSU rRNA sequences, we combined and optimized methods for producing full-length SSU rRNA cDNA from total RNA^4, 5^ with synthetic long read sequencing enabled by molecular tagging^6,7,8,9^. Full-length SSU rRNA molecules were enriched from extracted total RNA and converted to double-stranded cDNA, enabled by poly(A) tailing and single-stranded ligation, thereby avoiding the use of conventional SSU rRNA PCR primers and the resulting taxonomic bias^10^ (Fig. 1A). During first and second strand cDNA synthesis, the individual SSU rRNA molecules are uniquely tagged in both termini. The tagging enables preparation of short read sequencing libraries, where the resulting individual sequencing reads can be linked to the original template molecule. By sorting the short reads into separate bins based on their unique tag, full-length SSU rRNA molecules can afterwards be recreated using *de novo* assembly of the individual bins.

**Figure 1.**
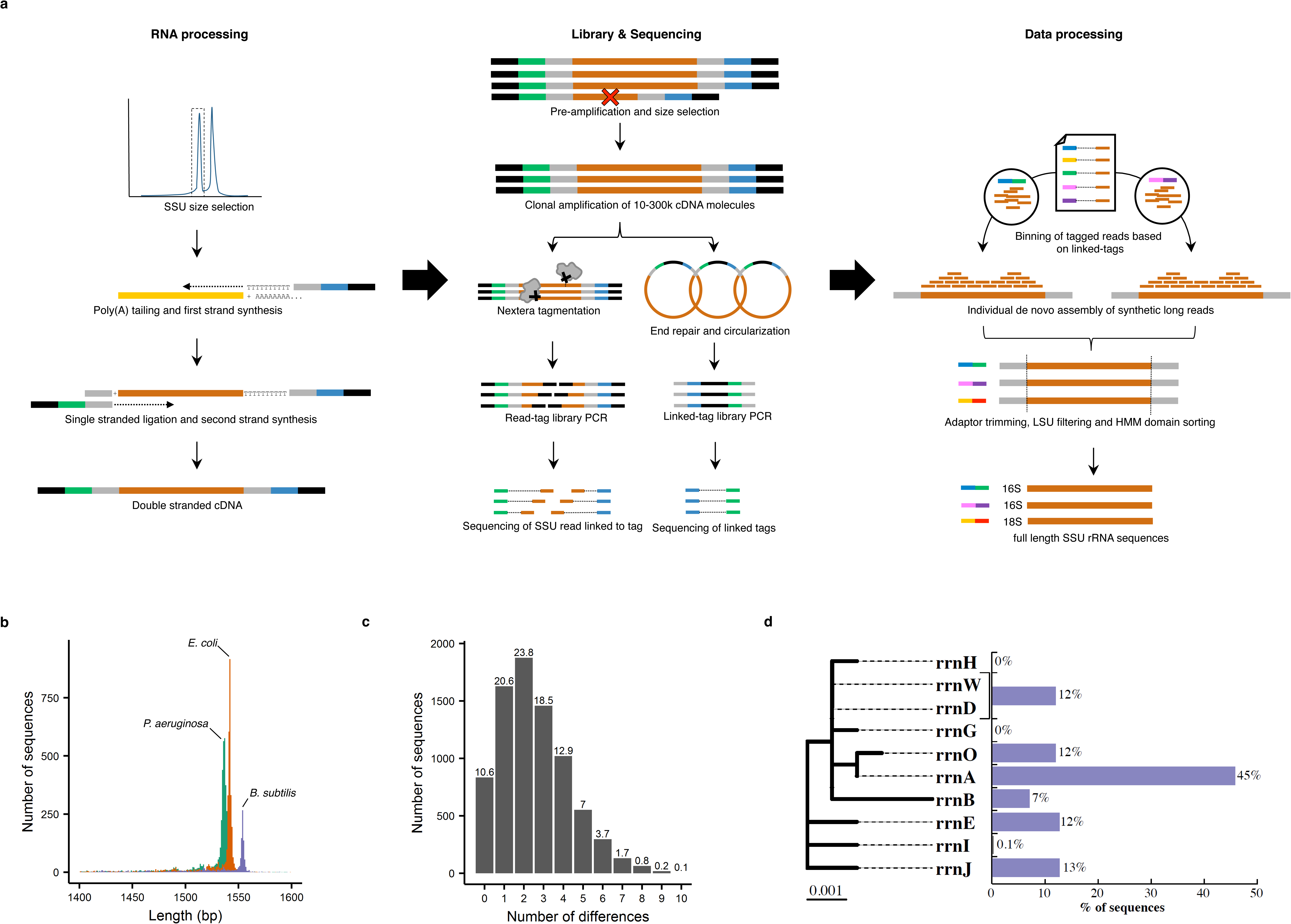
Overview and validation of full-length SSU rRNA sequencing. **a**, Schematic overview of the preparation of full-length SSU rRNA gene sequences from total community RNA (See **Fig. S1** for a detailed overview). First, SSU rRNA is enriched from extracted total community RNA using size selection. Then the SSU rRNA is polyadenylated, followed by reverse transcription and second strand synthesis. Adaptors used for first and second strand synthesis contain unique tags (green and blue), which in combination, become the unique “linked-tags” of the molecules. The cDNA is amplified with PCR and the product size selected to remove incomplete or truncated products. The full-length SSU rRNA amplicons are diluted to 10,000–300,000 molecules and amplified with PCR. The PCR product is split in two and used for preparing a read-tag library and a linked-tag library. The read-tag library is prepared by fragmenting the full-length SSU rRNA amplicons using Nextera tagmentation and library preparation. The resulting sequencing outcome is an internal SSU rRNA fragment read connected to a single unique tag read. The linked-tag library is prepared by circularizing full-length SSU rRNA amplicons to physically link the tags in close proximity. PCR is used to amplify the linked-tags, which are then identified with sequencing. The linked-tags are used to bin all SSU rRNA fragment tag-reads originating from the same parent molecule. Finally, *de novo* assembly is used to recreate the parent SSU rRNA sequence. **b**, Size distribution of assembled SSU rRNA sequences from the mock community. **c**, Error count distribution for raw SSU rRNA sequences from the mock community (Numbers indicate percent of all 16S rRNA sequences). **d**, The relative abundance of the different 16S rRNA genes for *B. subtilis*.

## Mock community evaluation

To estimate error and chimera rate of the method, we applied it to a mock community containing *E. coli* MG 1655, *B. subtilis* str 168, and *P. aeruginosa* PAO1, each with multiple 16S rRNA gene copies (4-10) that differ internally in 0 to 19 positions (up to 1.3% internal divergence). In a single Illumina MiSeq run, we generated 9,608 16S rRNA gene sequences over 1,200 bp (median 1,537 bp, Fig. 1B) with an average raw error rate of 0.17% (Fig. 1C) and a chimera rate of 0.19%. The raw error-rate corresponds well with the theoretical error-rate of the Taq DNA polymerase used in the PCR steps. Using standard error-correction, the average error-rate was reduced to 0.04%, with 62% of the sequences being perfect. The chimera-rate of 0.19% is up to 100 times lower than what can be observed in conventional PCR based studies^11^.

Even without error correction, the low error-rate enabled assignment of all full-length 16S rRNA sequences to their respective operons, exemplifying the resolving power of the method (Fig. 1D **and Fig. S2**). Interestingly, for *B. subtilis* three of the rRNA operons (rrn-I, rrn-H, and rrn-G) were not expressed. However, these are located closely together in the genome and also regulated by the same promoter^12^.

Earlier studies have indicated risk of taxa dependent biases in poly(A) tailing, due to modifications of the 3’-terminal ribonucleotide unit^4,5^, as well as biases from disruption of first strand synthesis due to internal modifications^13,14^. To investigate potential taxonomic bias, we compared full-length SSU rRNA sequences obtained from an activated sludge sample with total RNA shotgun sequencing of the same extracted RNA. All abundant taxa that were observed using shotgun RNA sequencing were also observed in the full-length sequences (**Fig. S3**).

## Error-correction of Oxford Nanopore data using molecular tagging

Tagging of individual molecules has been used as an effective consensus error-correction strategy in Illumina data^15,16^ and the principle is similar to the circular amplification strategies used to error-correct PacBio^17,18,19^ and Oxford Nanopore data^20^. Here we used the mock-community cDNA, designed for use on the Illumina MiSeq, and used it directly for Oxford Nanopore library preparation and MinION sequencing. Using uniquely tagged Nanopore reads and applying a naïve clustering and error-correction strategy, we increased the similarity from a median of 90% (range 69-97%) for the raw reads to a median of 99% for consensus reads generated from 7 or more tagged reads (range 98.7-99.6%, (**Fig. S4; Table S1**). With few additional adaptations, the molecular tagging approach can be optimized for use on the Oxford Nanopore platform, which should result in even lower error-rates, even for long DNA reads, currently not feasible for the circular amplification strategies.

## The method applied to real environmental samples

We used the full-length SSU rRNA approach to analyze samples from five widely studied ecosystems – soil, fresh water, human gut, anaerobic digestion (biogas production), and activated sludge (wastewater treatment). An average of 8685 rRNA sequences longer than 1,200 bp (median 1,434 bp) was obtained from each sample (**Table S2**). Each sequenced on a single Illumina MiSeq run. SSU rRNA made up 25-47% of all sequences, while large subunit (LSU) rRNA fragments made up the majority of the remaining sequences. The relative large fraction of LSU rRNA was unexpected, as the SSU rRNA peak was enriched using gel electrophoresis size selection (**Fig. S5**). However, LSU rRNA of many bacteria and lower eukaryotes also exist as nicked molecules, where one of the fragments has approximately the same size as the SSU rRNA^21,22^. In addition, degradation of stable RNA is more pronounced under conditions of starvation or environmental stress^14^. This is also in accordance with the experimental results obtained in this study, where more LSU rRNA was observed for the complex samples (53-75%) than for the mock community (8%).

We obtained SSU rRNA sequences from all domains of life, with representatives from 45 out of 66 bacterial phyla in the SILVA database^23^ including the majority of the known candidate phyla (Fig. 2A). To demonstrate that the method scales with sequencing capacity, we generated additional 62,140 rRNA sequences longer than 1,200 bp from the soil sample using a single Illumina HiSeq rapid run. From the single soil sample, we obtained 19,754 bacterial 16S rRNA sequences, which is equivalent to 18% of all soil-related sequences ever added to the databases^1^. Additionally, the 892 novel OTUs (97% clustering and > 3% difference to the SILVA database) obtained from the single soil sample, represent 8% of the new OTUs that are added to the SILVA database in a year^1^. For most environments, a single MiSeq sequencing run would add more eco-system specific sequences than ever added to the database for the particular environment.

**Figure 2.**
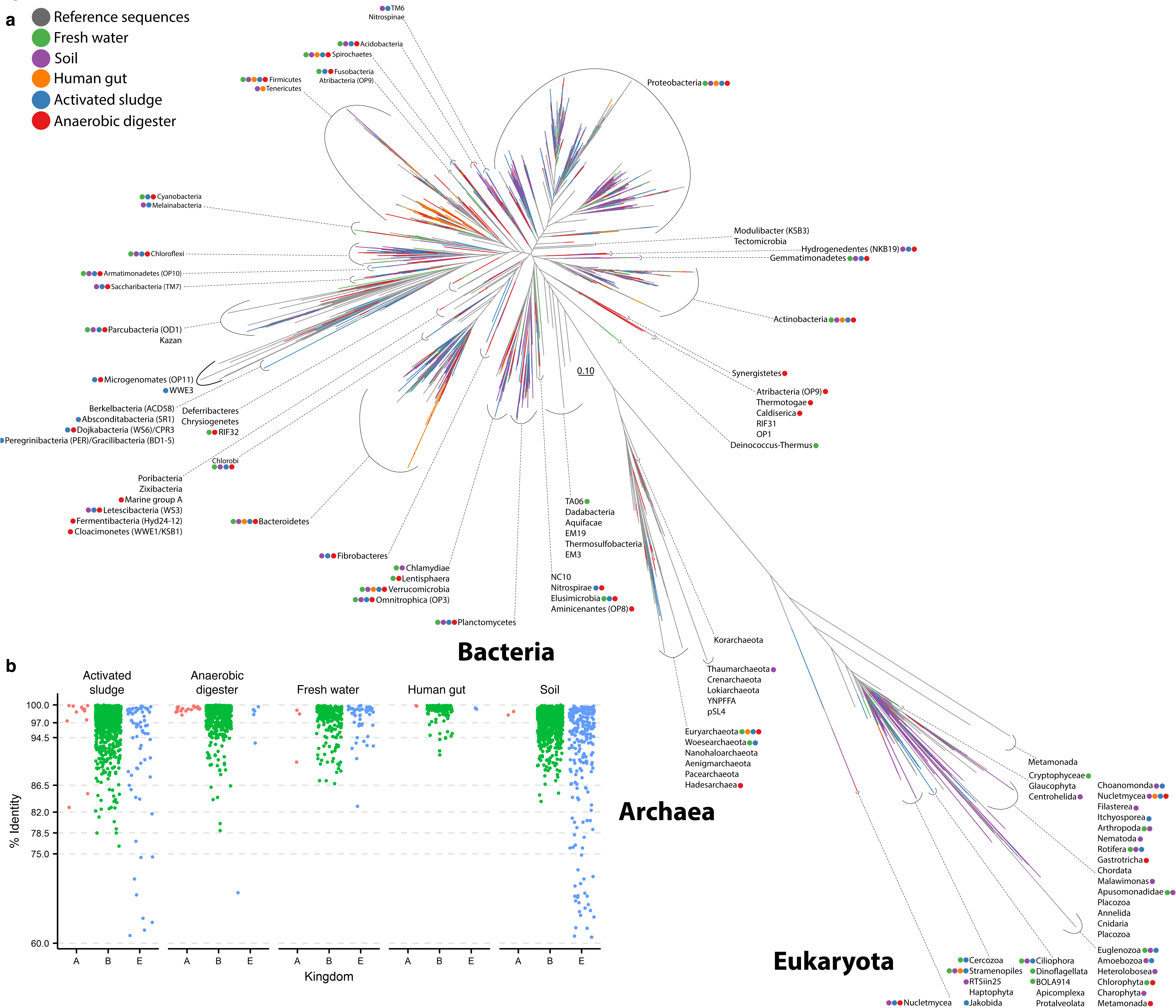
Coverage of the tree of life. **a**, Insertion of the newly generated SSU rRNA sequences to the current tree of life^31^. Brown branches represent sequences already in the public databases, and the other colors illustrate sequences added in this study. Note that the HiSeq soil data is not included. **b,** The percent identity of SSU rRNA gene sequences in the samples compared to their closest relatives in the SILVA database.

## Evaluation of bacterial diversity

Compared to the SILVA database, 30% of the full-length bacterial 16S rRNA OTUs represented new diversity (97% clustering and > 3% difference to the SILVA database). The degree of novelty was highly ecosystem specific. In the soil sample, 36% of the bacterial OTUs were novel compared to the database, while it was 5% in the human gut sample. These results underline that even in the densely sampled environments, as investigated in this study, a vast amount of bacterial diversity remains to be explored. We have refrained from attempting to define novel high-level phylogenetic groups based on our data, as it seems premature, when the databases will increase with orders of magnitude within a short timeframe. This will form a better foundation for robustly defining new phylogenetic groups.

A recent evaluation of primer bias using metagenomics estimated that up to 10% of bacterial diversity could be missed by conventional applied primers^2^. The generation of primer-free full-length 16S rRNA sequences in this study made it possible to access the conservation of the 27f and 1492r primers commonly used for generation of full-length sequences in the databases^24,25^. We found that 0 to 6% of full-length 16S rRNA OTUs had two or more mismatches to either the 27f or 1492r primer, depending on the environment (**Table S3**).

## Evaluation of eukaryotic diversity

In general, the eukaryotic 18S rRNA phylogeny is not well developed, especially not for the unicellular micro-eukaryotes. Universal eukaryotic primers have a poor coverage^26,27^ and they provide short amplicons with poor phylogenetic resolution^28,29^. To support this, we found a very high degree of novel eukaryotic diversity, when applying the primer-free approach. In total, 63% of the 18S rRNA OTUs were less than 97% similar to anything in the SILVA database (Fig. 2B), with 15% of all sequences being less than 80% similar to any known sequences. Recently, Hadziavdic *et. al*. (2014) developed a new set of universal primers for Eukaryotes, which target 76% of the SILVA database with perfect match and 93% with a single mismatch. Strikingly, when applied to the primer-free generated 18S rRNA sequences from this study, only 8% had perfect match to the primers and 80% had one mismatch (**Table S4**).

The new Eukaryotic Reference Database initiative (http://eukref.org/) has the goal to improve the eukaryotic reference databases. It is a collaborative annotation initiative to curate eukaryotic lineages by 18S rRNA gene data spanning the eukaryotic tree of life. Our full-length primer free approach will strongly support this endeavor and increase the power of high-throughput sequencing-based studies to discover fundamental patterns in microbial ecology.

## The beginning of a new era with a fully populated tree of life

The approach has fascinating perspectives in rapidly populating the tree of life. In this study alone, we have generated more than 30,000 full-length 16S rRNA gene sequences, which is approximately 15% of all sequences that were added to SILVA in 2015^1^. Our overall discovery rate of new diversity is higher than previously estimated based on the current databases^1^ and underlines that it is currently difficult to estimate the total bacterial diversity in the biosphere.

As the method is scalable and optimized to the most prevalent sequencing platform of today, we foresee a drastic increase in full-length SSU rRNA sequences that will be generated from all environments. It will be a monumental task to update the databases and difficult to maintain a phylogenetic tree encompassing all diversity. Our prediction is that ecosystem-specific databases, such as the human oral microbiome database^30^, will become more prevalent. Albeit decentralized, these databases might be easier to maintain and more information can be assigned to individual organisms based on the ecosystem context, which will make the databases more useful in practice.

It will be increasingly difficult to design both universal and specific primers. Instead, the high quality ecosystems-specific databases will be key to design new amplicon sequencing primers and fluorescence in situ hybridization (FISH) probes. For amplicon sequencing, this would mean better community coverage, compared to current universal primers. For FISH probes, it would be possible to design more specific probes, that increase the resolution of in situ single cell physiology studies, thereby aiding the task of linking identity and function in complex microbial communities.

In this study, we also recovered over 62,420 partial LSU rRNA fragments (1,200-1,600 bp). For comparison, there are 96,642 LSU rRNA sequences in the current release of the SILVA database (over 1,900 bp). Although the current implementation is limited to approximately 1,600 bp in order to maximize the yield of 16S rRNA sequences, a variation of the applied sequencing method has been demonstrated to yield multi-kb reads^9^. In addition, the promising error-correction of raw Nanopore reads demonstrated here is not limited by read length. Hence, also the LSU rRNA databases will experience a dramatic increase in the very near future.

The approach itself will allow researchers in microbiology and biology to get a complete community profile encompassing bacteria, archaea and eukaryotes, which has been difficult before. This would make it possible to look at interactions between the different domains of life in ecosystems, which have been scarcely studied until now.

## Acknowledgements

The study was funded by the Danish Research Council for Independent Research (FTP), the Innovation Fund Denmark, the Villum Foundation, and the Poul Due Jensen (Grundfos) Foundation We thank Holger Daims for insightful discussions of the manuscript.

